# Estimating glioblastoma migration-to-proliferation ratio from single time-point MRI data via radial density distribution

**DOI:** 10.64898/2026.10.26.761772

**Authors:** Rafael Bravo, David A. Hormuth, Ernesto A.B.F. Lima, Bahar Tercan, Boris Aguilar, Thomas E. Yankeelov

**Affiliations:** Oden Institute for Computational Engineering and Sciences, The University of Texas at Austin, Austin, TX 78712, USA; UT Austin Cancer Research Center, The University of Texas at Austin, Austin, TX, 78712; Department of Biomedical Engineering, The University of Texas at Austin, Austin, TX, 78712; Diagnostic Medicine, The University of Texas at Austin, Austin, TX, 78712; Texas Advanced Computing Center, Austin, TX, 78758; Institute for Systems Biology, Seattle, WA, 98109; Departme555.nt of Imaging Physics, The University of Texas M.D. Anderson Cancer Center, Houston, Texas, 77230

**Keywords:** Proliferation-Invasion model, reaction diffusion model, patient-specific parameter estimation, mathematical oncology, computational biology, magnetic resonance imaging (MRI), image-based modeling, prognostic biomarker

## Abstract

Glioblastoma (GBM) tumors with a high migration-to-proliferation ratio (*D*/*k*) are associated with treatment resistance, motivating the estimation of D/k from routine pre-treatment magnetic resonance imaging (MRI) data and the investigation of mechanisms underlying its heterogeneity across patients. Unlike previous approaches based on nested spherical approximations of enhancing and non-enhancing tumor volumes, we incorporate the spatial arrangement of the enhancing and non-enhancing tumor regions by deriving a 1D tumor density distribution from their segmentations and fitting a 1D reaction-diffusion model to estimate *D*/*k*. We achieved an error of under 5% when comparing our method to synthetic density distributions and found excellent correlation with synthetic tumor segmentations (Pearson correlation coefficient (PCC) = 0.996). We then evaluated associations between patient-specific *D*/*k* values and their clinical, transcriptomic, and radiomic features.

When applied to 102 pre-treatment MRI scans from the Brain Tumor Segmentation (BraTS) 2017 dataset, elevated *D/k* significantly correlated with reduced overall survival (log-rank *p* = 0.01), older age (*p* = 0.04), and a trend toward lower Karnofsky Performance Status (*p* = 0.059).

In a subset of 91 patients with available microarray data, mesenchymal-like phenotypes consistently exhibited elevated *D/k* across three established molecular subtype classifications (*p* = 0.013, *p* = 0.055, and *p* = 0.127). Elastic Net regularization followed by overrepresentation analysis revealed an enrichment of glycosylation-related pathways among genes associated with D/k variability (*p* = 0.0123 and p = 0.0368). Furthermore, Multi-Omic Factor Analysis identified a radiomic factor correlated with *D*/*k* (r = 0.57, p = 4×10^-9^), with the highest-weighted features corresponding to smoothness of the non-enhancing tumor region on T2 and FLAIR.

**Author Summary:** Glioblastoma tumors vary considerably in their growth patterns, with some forming relatively compact masses while others spread more diffusely into surrounding brain tissue. These patterns can be observed in a single pre-treatment MRI scan, where enhancing and non-enhancing regions approximate tumor cell density. Our method measures average tumor density in concentric spherical shells centered on the tumor core. It then fits a radially symmetric model of tumor growth to this distribution to estimate the ratio of tumor cell migration to proliferation (D/k).

We validated our method using simulated tumors with known migration and proliferation characteristics. We then applied our method to the publicly available BraTS 2017 dataset. We also used matched patient gene expression data to investigate molecular differences associated with variation in D/k.

We found that tumors with higher D/k were associated with poorer overall survival and a greater tendency toward the mesenchymal molecular subtype. We also identified an association between variation in D/k and genes involved in glycosylation-related pathways. Together, these findings connect tumor growth patterns visible on MRI with patient outcomes and molecular characteristics.

## Introduction

Pre-treatment magnetic resonance imaging (MRI) of glioblastoma (GBM) offers a window into unperturbed characteristics of these tumors. Several groups have investigated how this data can be used to infer clinically relevant tumor properties, including the tumor dynamics that produced the observed state, estimates of unseen tumor infiltration, and predictions of future growth^1–4^. One clinically important axis of interpatient variation is a tumor’s spatial distribution, which ranges from densely localized to highly diffuse^5,6^. This has been quantified as the ratio of effective tumor cell migration, *D*, to proliferation rate, *k*, denoted *D*/*k*, and can be estimated from a single pre-treatment MRI scan. *D*/*k* has demonstrated clinical significance, as high *D*/*k* is associated with IDH-mutant status^7^, reduced survival benefit from surgery^8^, poorer response to temozolomide chemotherapy^9^, and reduced overall survival^10^.

Current methods to estimate *D*/*k* are based on fitting a reaction-diffusion model to enhancing and non-enhancing tumor voxel volumes converted to nested spheres^7,11,12^. While the simplicity and consistency of these methods make them suitable for the clinical application, they cannot distinguish spiculated tumors from quasi-spherical tumors when both have similar numbers of enhancing and non-enhancing voxels, despite differences in their dispersal patterns. Irregular or spiculated tumors are associated with worse prognosis than quasi-spherical ones^13–15^, suggesting that incorporating this information could improve the clinical utility of *D*/*k*. Additionally, the cancer agent-based modeling literature suggests that irregular or spiculated growth patterns are associated with greater migration relative to proliferation^16–18^, linking this morphology to *D*/*k*.

In this study, we account for the additional migration required to form irregular, rather than spherical, tumor shapes. To better capture tumor spatial structure, we calculate a 1D radial density distribution from the tumor center and estimate *D*/*k* by fitting a reaction-diffusion model to this distribution. Improving *D/k* estimation may enhance treatment personalization by providing a more accurate characterization of patient-specific tumor invasiveness.

To evaluate the clinical and biological relevance of our approach, we apply it to the Brain Tumor Segmentation Challenge (BraTS) 2017 dataset, which contains pre-treatment tumor segmentations, demographic information, radiomic features, and outcomes for 102 GBM patients. This allows us to evaluate associations between *D*/*k* and overall survival as well as other clinical variables. Microarray data are available for 91 of these patients, facilitating the investigation of underlying relationships between gene expression, pathway enrichment, and *D*/*k*.

The balance between proliferation and migration is orchestrated by many overlapping and interacting pathways in GBM^19^. For example, deregulation of the PI3K/AKT/mTOR pathway through EGFR and PDGFR promotes proliferation^20^, and deregulation of Wnt and NF-κB pathways promotes migration^20^. There are also pathways and epigenetic factors that act as “switches” that modulate behavior, including the TGF-β pathway^20,21^ and the RAS/MAPK pathway^20^. These molecular differences are reflected in three major GBM transcriptional subtypes: Classical, Proneural, and Mesenchymal^22^. Classical tumors are characterized as EGFR-associated and proliferative, Proneural tumors as PDGFRA-driven and progenitor-like, and Mesenchymal tumors as NF-κB mediated, migratory, and drug resistant^23^. Because patient tumors in our dataset have been classified using several molecular subtype classifications^24,25^, we can test for associations between these subtypes and *D*/*k*.

Building on these observations, we investigated how interpatient differences in gene expression are associated with inferred differences in proliferation and migration through the lens of *D*/*k.* We used Elastic Net modeling and gene-set variation analysis to identify genes and pathways that are associated with variation in *D*/*k*. We integratively analyzed the radiomic and the microarray data using Multi-Omics factor analysis (MOFA)^26^ to identify latent factors associated with D/k. Recent evidence suggests that *D*/*k* may reflect patient-specific molecular heterogeneity: Metz et al. found that *D*/*k* correlated with inferred molecular activity related to proliferation, apoptosis, and cell-cell adhesion^10^.

*D*/*k* is estimated from a model of tumor growth and migration, thereby providing us with an interpretable link between tumor behavior at the whole-tumor scale (from imaging-based modeling) and the underlying processes at the cellular and molecular scale (from gene expression analysis). To investigate these relationships, we develop and validate a pipeline for estimating *D*/*k* from pre-treatment MRI and apply it to the BraTS 2017 cohort. Using the clinical, molecular, and radiomic features from this dataset, we examined associations between *D*/*k* and patient outcomes, established glioblastoma subtypes, gene expression, and multi-omic factors.

## Methods

### Pipeline Overview

Figure 1 provides an overview of the *D*/*k* estimation pipeline and BraTS dataset analysis. We first developed the pipeline for computing *D*/*k* and evaluated its accuracy using synthetic tumor density distributions and tumor segmentations. We then calculated *D*/*k* for 102 pre-treatment tumors from the BraTS 2017 dataset, performed Kaplan-Meier analysis, and evaluated correlations between *D*/*k* and clinical variables. Finally, for 91 of the BraTS 2017 patients with available microarray data, we employed three techniques to link gene expression to *D*/*k*: (1) assessing correlation between *D*/*k* and previously defined GBM transcriptional subtypes; (2) using Elastic Net regression with leave-one-out cross validation (LOOCV) to identify *D*/*k-*predictive genes, followed by overrepresentation analysis (ORA) to determine enriched pathways; and (3) applying MOFA, to project gene expression and radiomic features onto a low-dimensional set of factors to test for correlation with *D*/*k*.

**Figure 1:**
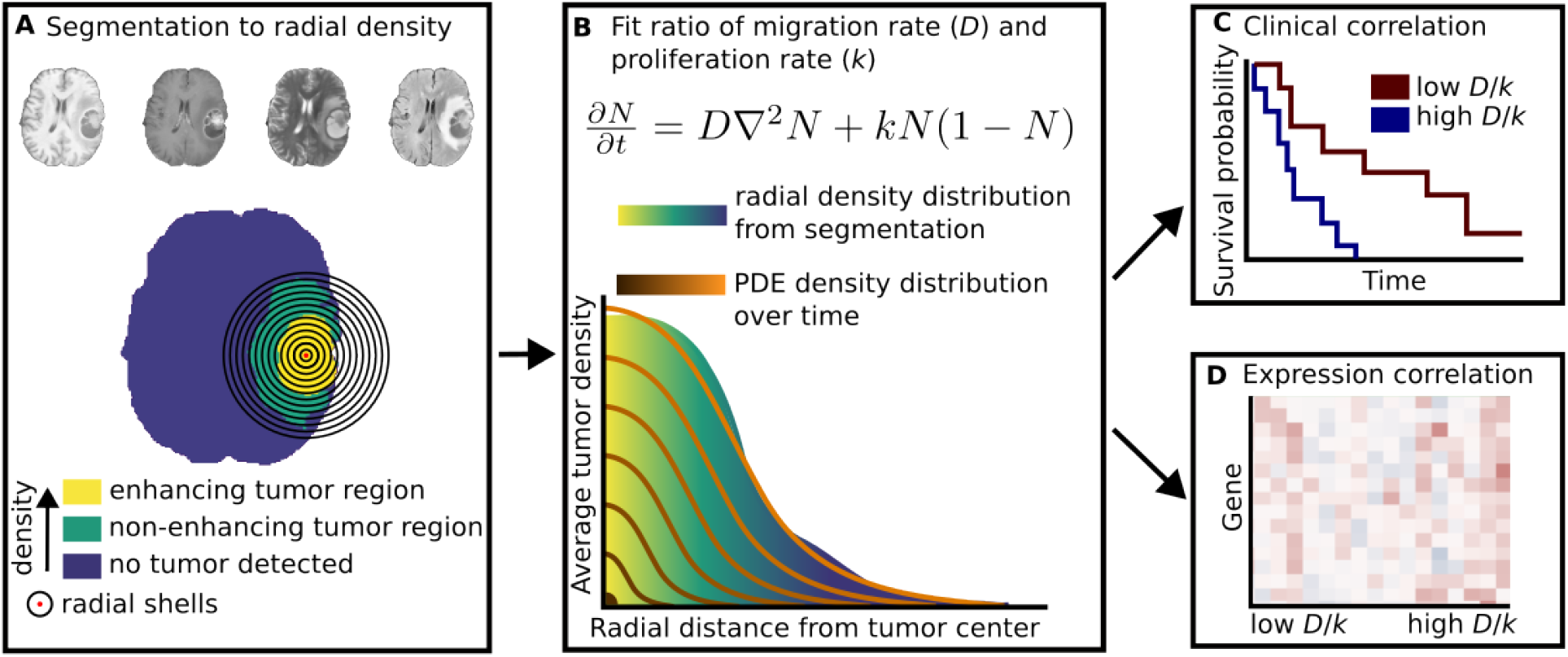
Overview of the D/k estimation and multi-modal analysis pipeline. **(A)** From left to right, pre- and post-contrast *T_1_*-weighted, *T_2_*-weighted, and FLAIR MRI images are segmented into enhancing and non-enhancing tumor regions. The center of mass of the largest enhancing region (red dot) is defined as the tumor initial position, and brain voxels are binned into spherical shells based on their distances in millimeters to this tumor initial position (black rings). **(B)** Average tumor densities within each shell yield a one-dimensional radial density distribution. A one-dimensional reaction-diffusion model (Equations 2–4) is fitted to this density distribution, and the *D*/*k* ratio minimizing the sum squared error was identified. **(C)** Estimated D/k values were evaluated for clinical relevance via Kaplan-Meier survival analysis and clinical feature correlations **(D)** Bulk RNA microarray expression profiles were correlated with *D*/*k* to identify associated genes and pathways.

### BraTS 2017 dataset: imaging, radiomic, and microarray features

We used the publicly available BraTS 2017 dataset^27–29^, comprising pre-treatment segmented MRI scans from 102 GBM patients, with matched RNA microarray data available for 91 individuals. Patient demographics are detailed in Table S1.

The dataset includes skull-stripped pre- and post-contrast *T_1_*-weighted, *T_2_*-weighted, and *T_2_* Fluid-Attenuated Inversion Recovery (FLAIR) images (Figure 1A). We grouped the tissue into three types based on the manual segmentations provided with the dataset^28^: peritumoral edema (which we considered the non-enhancing tumor region), the combined enhancing tumor and necrotic/non-enhancing tumor core (which we considered the enhancing tumor region), and the healthy-appearing brain (which we considered the normal tissue). All quantitative analyses were performed using the brain mask and the tumor segmentations. All scans and segmentations were provided with 1 mm^3^ spatial resolution. Details of the MRI acquisition and segmentation can be found in refs. ^27^ and ^28^, respectively.

Gene expression data (mRNA microarray) were available for 91 of the BraTS 2017 patients, who are also part of The Cancer Genome Atlas (TCGA)^30^. We downloaded this microarray data from FireBrowse^31^. The data was Robust Multi-array Average (RMA) normalized and we used it for selecting predictive genes for *D*/*k* via Elastic Net regression, gene set enrichment, and MOFA analyses.

The BraTS 2017 dataset also provides 704 radiomic imaging features extracted from the pre-treatment MRI sequences and tumor segmentations for each patient, including measures of tumor intensity, shape, and texture, generated by Bakas et al.^28^. We integrated these features with the microarray data in our MOFA analysis to identify factors capturing shared variation across imaging and gene expression that correlate with *D*/*k*.

### Mapping MRI data to 3D tumor density distribution

To estimate *D*/*k* from segmented MRI data, we first approximated each patient’s 3D spatial tumor density distribution, then generated a 1D radial density distribution from this 3D distribution and subsequently fitted our PDE to the 1D distribution.

Although apparent diffusion coefficient (ADC) values can estimate cell density^32–34^, these are unavailable in the BraTS 2017 dataset. We therefore assigned an approximate tumor density value for each voxel based on its region: enhancing tumor, non-enhancing tumor, or normal tissue. Previous biopsy studies guided our assignment of density values to voxels within the enhancing, non-enhancing, and normal-appearing segmentations^35^. To simplify the model and focus on tumor migration and proliferation, we did not model necrosis.

We assumed that the edge of the enhancing region is approximately 80% as dense as the non-necrotic tumor center (supported by biopsies^35^ and consistent with prior GBM modeling studies^11^) and inferring that the average density in the enhancing region falls between 80% and the maximum capacity (100%) of the tumor center. We thus assigned a fixed, average density of 90% throughout the enhancing region. We set the density in the non-enhancing region at 45% of the carrying capacity based on image-guided biopsy studies that indicate tumor cell density in the non-enhancing region is approximately half of the density in the enhancing region^36–39^. For the healthy-appearing brain outside the enhancing and non-enhancing regions, we set these voxels to have zero tumor cells. By assigning these three tumor cell density values as a proportion of the carrying capacity (i.e., 90% in enhancing, 45% in non-enhancing, and 0% in healthy-appearing brain), the tumor segmentations were transformed into 3D density distributions.

### Mapping 3D tumor density distributions to 1D radial density distributions

To fit our 1D PDE, we generated a 1D radial density distribution from the 3D voxel-level density distributions. We defined the tumor initial position as the center of mass of the largest contiguous enhancing region (Figure 1A, red dot). We binned all voxels within the brain mask based on their distance to the tumor initial position in 1-mm-thick spherical shells (Figure 1A, black rings). The voxel closest to the initial position was always placed in the first bin, voxels whose centers were between 0.5 mm and 1.5 mm of the initial position were placed in the second bin, and all voxels whose centers were between 1.5 mm and 2.5 mm of the initial position were placed in the third bin. This binning process continued up to 99.5 mm, for 100 total bins. To get a 1D density distribution from these bins, we averaged the densities of all voxels in each bin, (Figure 1B, yellow-green-blue color gradient).

### Mathematical model

The PDE that we calibrated to each patient’s 1D density distribution is based on the reaction-diffusion equation:

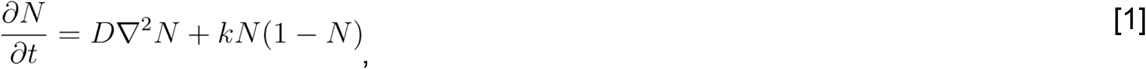

where *N* is the tumor cell density in space and time, *D* is the (spatially invariant) tumor cell diffusion coefficient (mm^2^/time) and *k* is the (spatially invariant) tumor cell proliferation rate (1/time). As our model consists of isotropic diffusion (*D*) and proliferation (*k*), we use a spherical 1D PDE to fit the spatial density distribution, thereby reducing the simulation complexity while preserving the capacity to represent tumor spread.

To solve Eq. [1], we use a standard 1D finite volume implementation of a 3D radially symmetric system^40^ as described in Eqs. [2] – [4]:

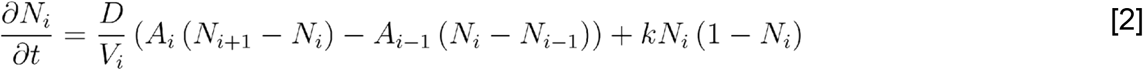

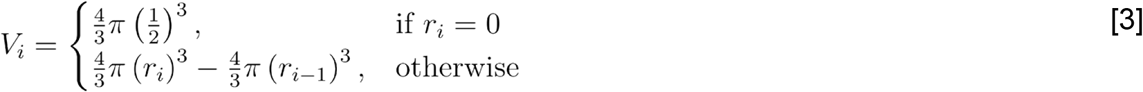

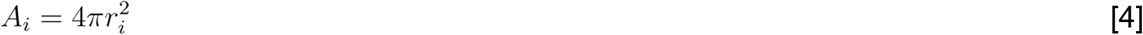

where *N_i_* is the average tumor cell density within radial shell *i*, *V_i_* is the volume of radial shell *i* (Eq. [3]), and *A_i_* is the area of the interface between radial shell *i* and radial shell *i* + 1 (Eq. [4]), *r_i_* is the radial distance to the outer surface of radial shell *i*.

The finite volume approach accounts for the volume of each spherical shell as in Eq. [3] as well as the fluxes between spherical shells (which are proportional to the area of the interface between adjacent shells (Eq. [4])). We used the finite volume implementation because it is flux/mass conserving, handles volumes defined in spherical coordinates directly, and behaves robustly at the center voxel where naïve finite differences implementations can encounter difficulties^41^. To initialize Eq. [2], we assume an empty domain with a single tumor cell located at the tumor’s initial position (*I* = 0). If we assume 1e6 cells/mm^3^ as the tumor cell density at carrying capacity^42^, then the initial normalized density in the central voxel is 1e-6. We assume zero flux boundary conditions.

### D/k estimation via model fitting to patient data

The fitting is accomplished in two stages: (1) a grid search finds a global estimate of the combination of diffusion rate *D* (mm^2^ per model timestep), and a simulation duration in timesteps *t* (unitless), that minimizes the squared difference between the density distributions from the data and the model. (2) the Levenberg-Marquardt optimization algorithm^43^ is initialized with the result of the first stage and iterates until convergence.

The time required for the tumor to reach the state observed at diagnosis cannot be inferred from a single MRI timepoint, as at least two timepoints are required to calculate change over time. Therefore, simulation timesteps, *t*, represent iterations of the PDE, not a real-world duration. Also, we hold the proliferation rate *k* constant throughout fitting because we are interested in the ratio *D*/*k*, and unique values of *D* and *k* cannot be estimated from just one observation.

During fitting, we weighted the error of each shell proportionally to its volume (Eq. [3]). Weighting by shell volume as opposed to voxel counts ensures that the fitting objective does not vary between patients due to differences in brain boundary truncation, which is consistent with our spherically symmetric representation of the tumor.

The grid search was conducted by first generating a grid of tumor density distributions with different diffusion rates (*D*) and different model durations (*t*) that could be compared to the target density distribution. This grid was generated with the proliferation rate (*k*) fixed at 10^-4^ per timestep and *D* ranging from 1E-5, to 1E-2, with 1000 log-scale increments (meaning *D/k* varied from 1E-1 to 1E2, which covers the range that we see in the patient data fits, see Figure S1), and a range from 0 to 1E6 timesteps with 1000 equally spaced increments. This generated a 1000 × 1000 grid of density distributions. For each distribution in the grid, we calculated the sum squared error between the entry and the patient-derived density distribution.

The Levenberg-Marquardt algorithm was used to improve the fit after the grid search, using the same bounds as the grid search. Levenberg-Marquardt was run until any one of the following criteria was met: 100 iterations elapsed, the sum squared error did not improve for 25 consecutive iterations, or after 10 improving steps, the sum of squared errors was still within 1E-4 of the value before those improvements.

Figure 2 visualizes the model fitting process for three representative patients that span the range of *D*/*k* values found in the BraTS 2017 dataset. We first calculated the tumor initial position from the center of mass of the enhancing region and found the average tumor density in radial shells (Figure 2A, B). We next used a grid search and Levenberg-Marquardt optimization to find the best fitting *D*/*k* and *t* for the patient (Figure 2C, D). We also show how well the fits match the density distributions generated from the patient data (Figure 2E).

**Figure 2:**
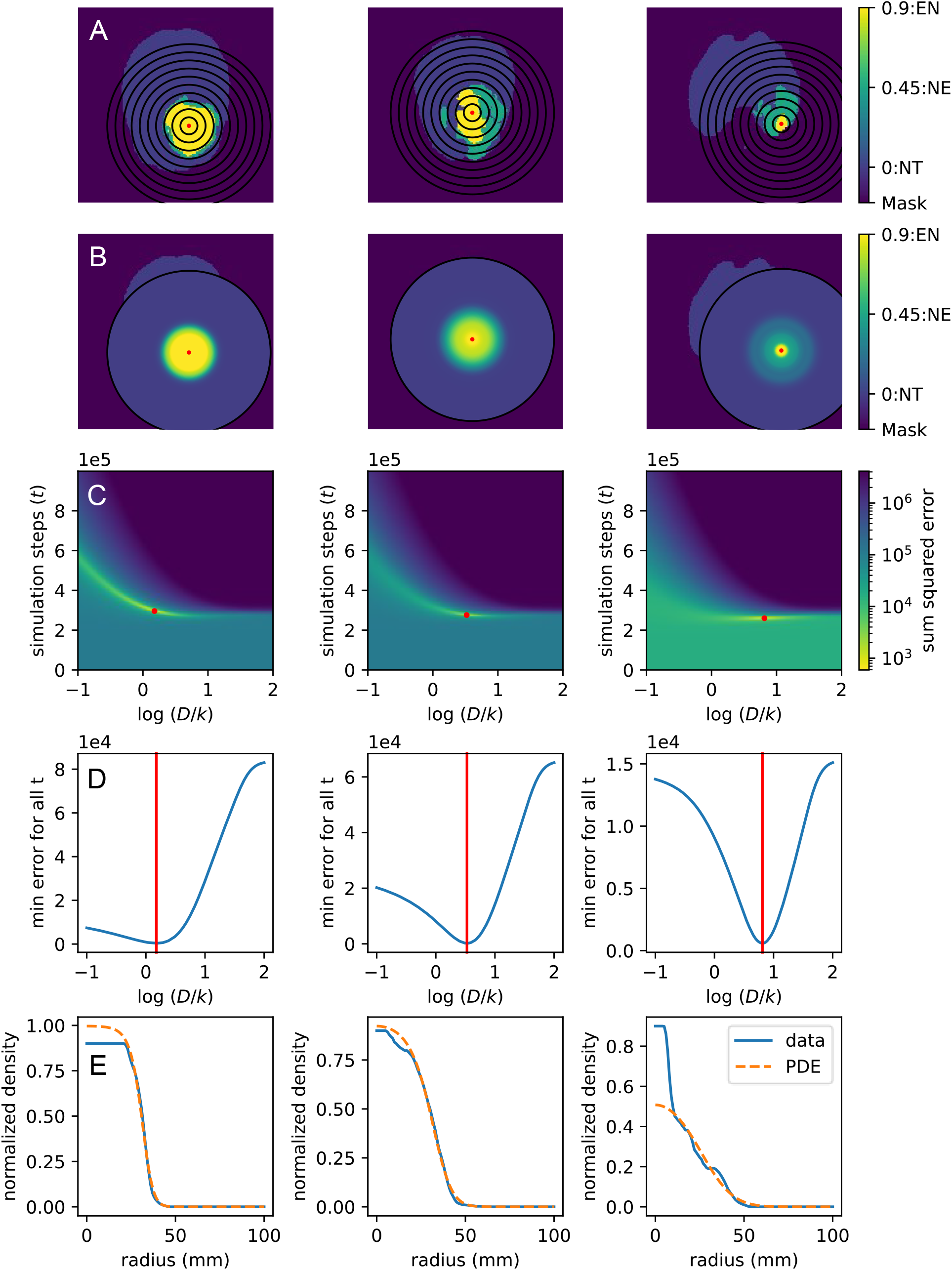
Representative images of the *D*/*k* algorithm are shown for three example patients from the BraTS 2017 dataset, one in each column. A A single slice from the 3D MRI segmentation is shown. First, the center of mass of the enhancing region is found (red dot), then the voxels are binned based on their distance to the center (black rings highlight every 10 mm to the center) **B** The radially symmetric density distribution is overlaid onto the MRI slice to visualize the conversion from 3D spatial data into a 1D density distribution. **C** Heatmaps show sum squared error of all parameter combinations used in grid search for example patients. The red dot indicates the minimum error point, which is used to initialize the Levenberg-Marquardt algorithm to provide estimates of *D*/*k* and *t* for each patient. **D** The minimum sum squared error for each *D*/*k* ratio is shown (blue line), summarizing the results of the grid search in C. The vertical red line shows the lowest error *D*/*k* ratio that was used as an initial position for Levenberg-Marquardt. **E** The blue line shows the 1D density distribution derived from the patient segmentation data, the orange dashed line shows the density distribution from the best-fitting patient-specific *D*/*k* ratio. Plots in the other two columns show the same information as A-E but when the algorithm is applied to other patients.

All steps of the PDE, grid search, and Levenberg-Marquardt algorithm were implemented in Python. The ‘numba’ library was used to accelerate PDE computation. Simulations were run on a Macbook Air with a 10-core Apple M4 Chip with 16GB of RAM.

### Simulated density distributions

We generated synthetic tumor density distributions to test whether our approach could extract the correct *D*/*k* when it is known. We first tested this with a 3D finite difference forward Euler solve of Eq. [1]. The PDE initial condition was an empty 199 X 199 X 199 mm^3^ domain with a single-cell density (1E-6) in the center voxel. The system was then run forward in time until the integrated tumor density reached up to 1/10th of the total domain volume. This process was repeated with *k* fixed at 1E-2, and *D* ranging from 1E-3 to 1E-1 in 10 log-scale increments; thus *D*/*k* ranged from 1E-1 to 1E1. To generate a 1D density distribution from these 3D domains, we computed a line profile starting with the center voxel and moving along the positive *x*-direction to the edge of the domain. The 100 voxels selected in this way became the 1D density distribution that we used for testing.

To test how varying spatial scales would impact the accuracy of the *D*/*k* fit, we also generated 3D density distributions with decreased spatial resolutions. These used the same size domains, but with larger volumes per voxel and scaling the starting density in the center-voxel to account for the volume of a single cell in a larger voxel.

We also added noise to the line profiles to test how it would alter the accuracy of the *D*/*k* estimation. We added noise by sampling a normal distribution for each voxel, adding these values to the original voxels, then bounding the results between 0 (empty voxel) and 1 (the voxel carrying capacity). We generated 20 random replicates in this way per *D*/*k* value and with different noise standard deviations (σ = 0.01, 0.05, 0.1). For reference, Grech-Sollars et al.^44^ reported mean coefficients of variation of 2.3% across repeated measurements of diffusion MRI parameters in healthy brain tissue, corresponding to roughly in between our σ = 0.01 and σ = 0.05 scenarios.

### Simulated tumor segmentations

We also tested whether the correct *D*/*k* would be estimated if the synthetic density distributions were first segmented into enhancing, non-enhancing, and healthy-appearing regions. To test this, we started with the same forward Euler method for generating 3D synthetic density distributions that we used for the previous test. We then assigned densities ≥ 80% as enhancing tumor, densities between 80% and 10% as non-enhancing tumor (with thresholds chosen symmetrically around the literature-derived average non-enhancing tumor density of 45%), and densities ≤ 10% as healthy-appearing brain. We then converted these three regions into our literature-derived average densities of 90%, 45%, and 0%, respectively and ran our algorithm to estimate *D*/*k* on the 3D synthetic regions. Specifically, we found the center of mass of the enhancing region, binned the voxels by their distance to the center of mass, averaged the density in these bins to derive a 1D density distribution, and fitted our 1D PDE to it to estimate *D*/*k*.

With this segmentation method, we tested how the *D*/*k* estimations would differ if the synthetic tumors were segmented after growing to different sizes, and if the tumors were close to the edge of the domain, such that the boundary would cause asymmetric growth.

### Clinical Data and Survival Analysis

We used clinical information including subtypes, age, Karnofsky Performance Score (KPS), OS time and OS status from ^25^. For survival analysis, we used the lifelines python package for plotting KM plot and surv_cutpoint function from survminer R package to find the cut point for D/k to segregate poor and favorable survival groups. The reported p-value was not adjusted for cutoff selection.

### Elastic Net analysis

We applied an Elastic Net approach to identify genes whose expression levels are predictive of *D*/*k*. This required tuning four hyperparameters. To limit the number of input features to Elastic Net, we tuned a variance threshold (hyperparameter 1) to remove genes with low gene expression variance. A correlation threshold was also tuned (hyperparameter 2) to keep only a subset of genes that were the most correlated or anti-correlated with *D*/*k*. For Elastic Net, we also determined the weighting of the ratio between Least Absolute Shrinkage and Selection Operator (LASSO)^45^ and ridge regression^46^ (hyperparameter 3), and the weighting between these penalties and least-squares (hyperparameter 4). We used leave-one-out cross-validation (LOOCV) to test different combinations of these four hyperparameters to find the combination that maximized the PCC when predicting the *D*/*k* values of the left-out patients.

With the resulting set of genes found by Elastic Net using the optimal hyperparameter combination, we performed overrepresentation analysis (ORA)^47^ using the Reactome and MsigDB Hallmark gene sets to find pathways that relate to the most predictive genes.

### Multi-omics factor analysis (MOFA)

We applied MOFA^26^ to identify latent factors derived jointly from the microarray expression data and radiomic features. MOFA is an unsupervised dimensionality reduction method that identifies latent factors representing the principal sources of variation from multi-modal datasets. It also distinguishes factors shared across different data modalities from those exclusive to individual ones. We then tested for correlations between *D*/*k* and the resulting factors.

## Results

### Validating D/k fitting using simulated density distributions

We first determined if our calibration approach (i.e., a grid search to initialize the Levenberg-Marquardt algorithm) could accurately estimate the *D*/*k* ratio from line profiles taken through a 3D object simulated using Eq. [1] (Figure 3A, B). Since we are fitting a ratio, we can either hold *D* constant and fit *k* or vice versa. To decide which approach to take, we compared how accurately each approach recovered the known *D*/*k* ratios from the line profiles (Figure 3C) and the number of timesteps required for fitting (Figure 3D). Fixing *k* or fixing *D* yielded concordance correlation coefficients (CCC) of 0.9979 and 0.9978, respectively. However, as the interquartile range of the number of timesteps used for fitting was smaller with *k* held constant (median 4.4 × 10^5^, IQR 3.36 × 10^5^–6.35 × 10^5^), than with *D* constant (median 4.37 × 10^5^, IQR 1.99 × 10^5^–1.05 × 10^6^), we decided to hold *k* constant and fit *D*.

**Figure 3:**
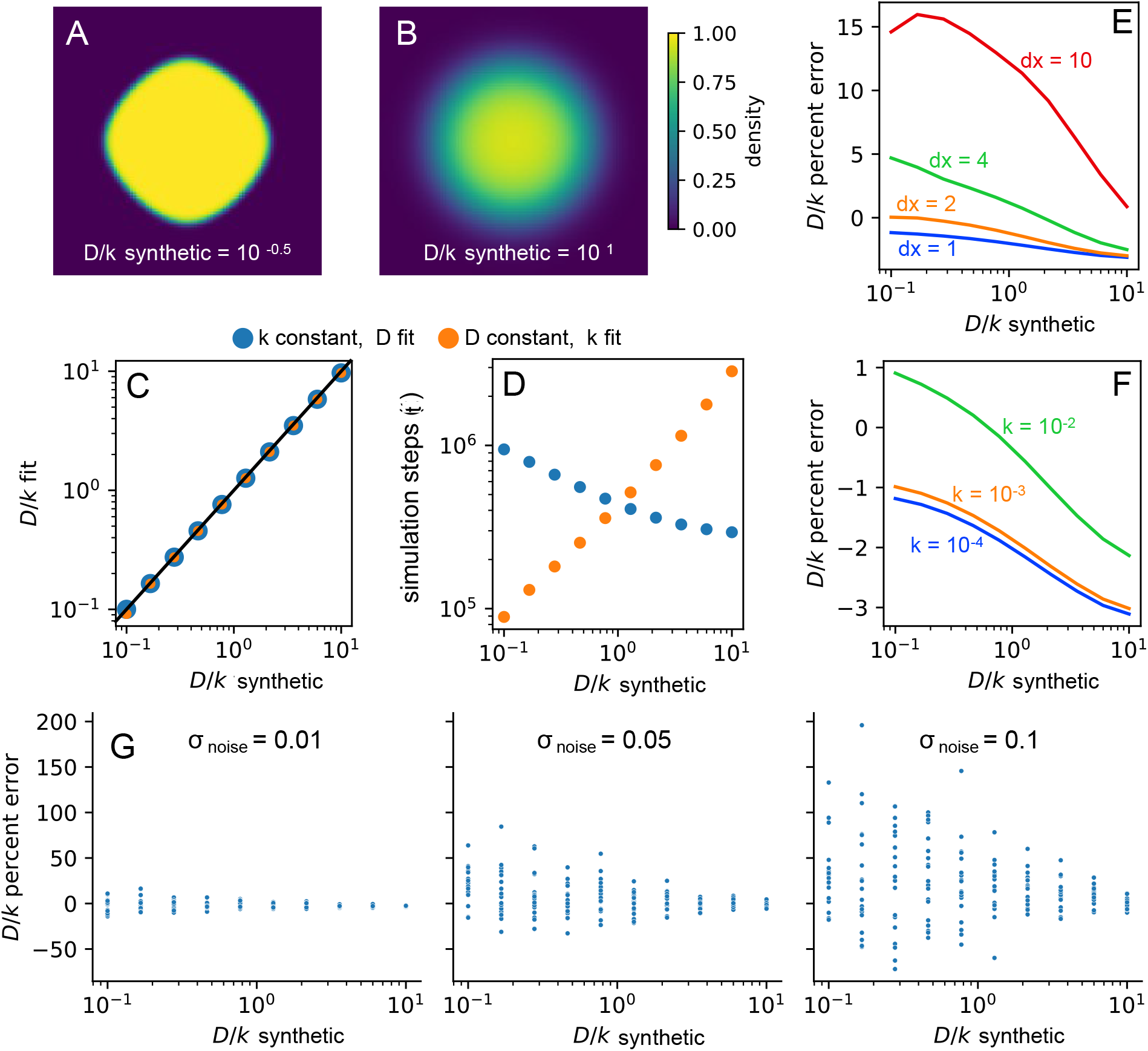
Validation using line profiles from synthetic density distributions. A, **B** Example synthetic 3D density distributions. These distributions were generated by running the model in an initially empty 3D domain (199 x 199 x 199 mm^3^), starting with the single cell initial condition in the center voxel, and stopping when the domain is 10% filled. **C** *D*/*k* values were derived with *k* fixed and *D* fixed, respectively, on 10 logarithmically spaced synthetic density distributions (*D*/*k*_synthetic_). **D** number of model timesteps needed to converge on the best fitting *D*/*k* values. **E** Percent difference between *D*/*k* fitting result and synthetic density distributions with different spatial resolutions (*D*/*k* true). **F** Percent difference between the estimated *D*/*k* value and *D*/*k* from the synthetic density distributions with different temporal resolutions (*k* value per unitless timestep). **G** *D*/*k* percent differences for 20 repeats of the *D*/*k* fitting method run on synthetic density distributions after the addition of Gaussian noise with the specified standard deviation.

We next tested the impact of different spatial scales (by changing grid resolution, Figure 3E) and different temporal scales (by changing the fixed proliferation rate, Figure 3F) on the fitting error. We observed that our approach estimated *D*/*k* with a median percent error of -2.01% (IQR -2.66% to -1.48%) at a spatial resolution of *dx* = 1 mm, and the temporal resolution set by *k* = 10^-4^ per model timestep. Finally, we introduced Gaussian noise to the line profiles to determine how this impacts parameter estimation (Figure 3G). The noise caused greater deviations in the estimation of *D*/*k* for small *D*/*k* values (e.g. 0.1 mm^2^) than for large *D*/*k* values (e.g. 10 mm^2^). For example, when σ_noise_ = 0.05 and *D*/*k_synthetic_ = 0.1* mm^2^, the mean percent error (standard deviation) was 20.1% (20.5%), while for *D*/*k*_synthetic_ = 10 mm^2^, the mean percent error was -1.2% (2.7%).

### Validating the D/k pipeline using simulated tumor segmentations

We determined if we could accurately fit *D*/*k* when the synthetic density distributions were first separated into simulated tumor segmentations analogous to real enhancing and non-enhancing segmentations (Figure 4A). This involved finding the correct initial position of the tumor, generating a 1D density distribution around that initial position, and calibrating Eq. [2] to that distribution with the grid search and Levenberg-Marquardt. We also wanted to check the accuracy of the *D*/*k* estimation across tumors of different sizes. We plotted the resulting fits across different permutations of *D*/*k* and tumor size to determine whether the resulting *D*/*k* fits would align with the original synthetic *D*/*k* values (Figure 4B). We found that the estimates of *D*/*k* were offset from the synthetic *D*/*k* values (*CCC = 0.813*), though they were highly correlated (*PCC = 0.996*). We also explored the impact of initiating the tumor near the edge of the domain. The results from this test were provided in Figure S2.

**Figure 4:**
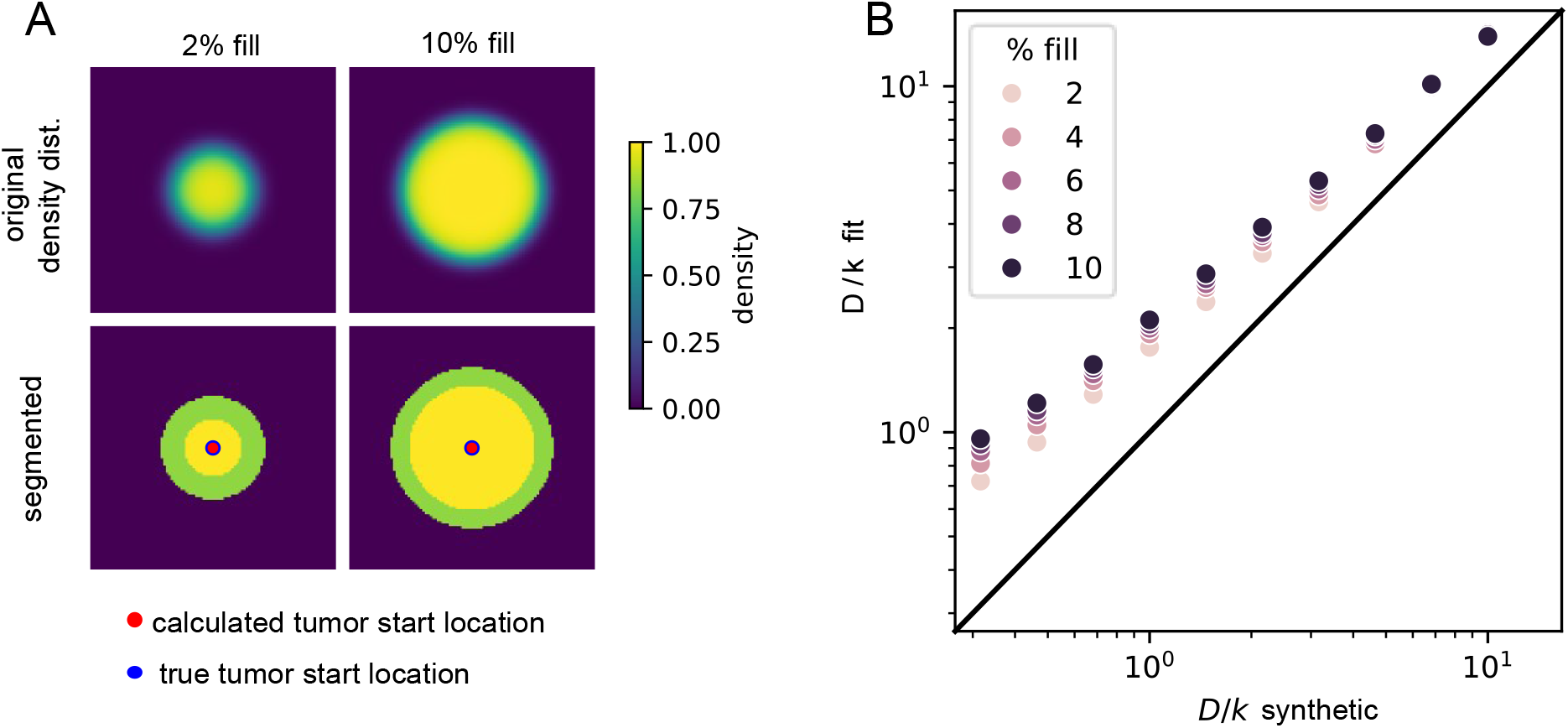
Validation using synthetic tumor segmentations. **A** Synthetic density distributions and the results of enhancing (solid yellow) and non-enhancing segmentations (solid green). Tumor initial location (red dot) was calculated from the segmentation using the center of mass of the enhancing region. Percent fill indicates total tumor density at the time of segmentation as a fraction of the maximum carrying capacity of the domain if fully saturated. **B** *D*/*k* calculated via the *D*/*k* algorithm compared to the synthetic data *D*/*k* with different fill proportions.

### Correlation with clinical variables and molecular subtypes

After estimating *D*/*k* for all 102 patients from the BraTS 2017 dataset, we investigated whether *D*/*k* correlated with clinical variables. To test whether *D*/*k* is associated with overall survival, we performed a log-rank test, sweeping the threshold from the 10th to the 90th percentile, to maximize separation in overall survival between high and low *D*/*k* groups (threshold = 2.081 mm^2^; Figure 5A). Patients in the high *D*/*k* group (n = 54) showed significantly worse overall survival than patients in the low *D*/*k* group (n = 47) (p = 9.9X10^-3^). Survival data were unavailable for one patient. Age at diagnosis was significantly correlated with *D*/*k* (p = 0.04; Figure 5B), and Karnofsky Performance Score (KPS) showed a noticeable trend (p = 0.06; Figure 5C). We conducted these same analyses using *D*/*⍴* as defined in Swanson et al. ^11^ and found similar trends (Figure S3). We also tested whether *D*/*k* was significantly associated with molecular subtypes by using the subtypes originally identified in^24^ from microarray analysis of a larger GBM dataset that included the BraTS patients. We compared the mesenchymal subtype to all other subtypes combined; the neural subtype was excluded as it is thought to reflect contamination with normal cells^22^. The mesenchymal subtype showed a higher mean *D*/*k* relative to the remaining subtypes, nearing statistical significance (p = 0.0547) (Figure 5D). Similar trends were observed using the Transcriptome subtype, (p = 0.127) and Random Forest Sturm cluster subtype, (p = 0.0133) both from^25^ (Figure 5E,F).

**Figure 5:**
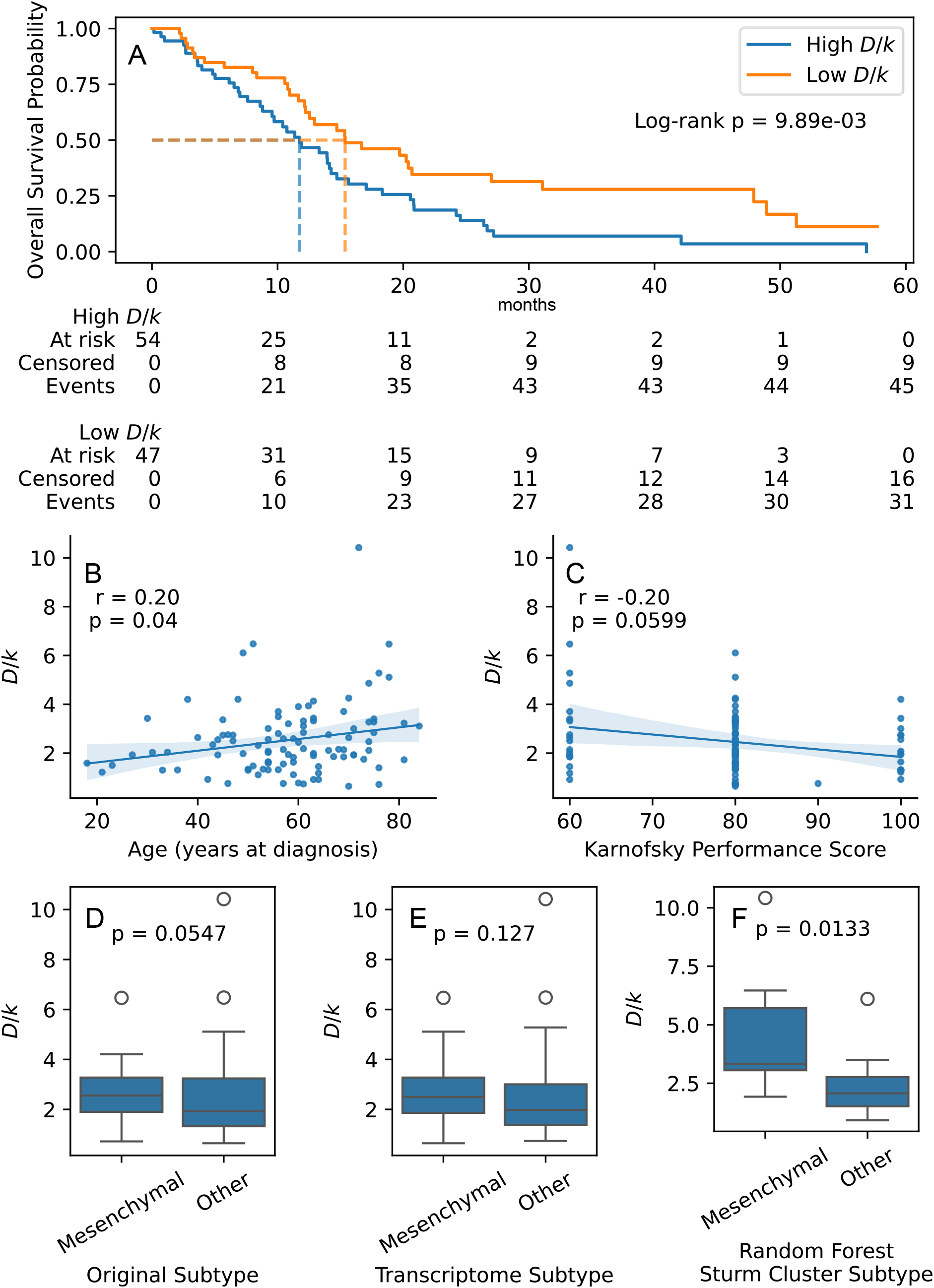
Association of D/k with patient survival, clinical characteristics, and molecular subtype. **A** Kaplan-Meier curves of optimally separated groups (High and Low *D/k*) are shown with associated log-rank p-value. Dashed lines show median survival. **B, C** Correlation of patient age with *D*/*k* and Karnofsky Performance Score with *D*/*k*. Spearman correlation was used to calculate r and p values. **D-F** Mesenchymal clusters compared to all other clusters combined; p-values were computed using Mann-Whitney U test. **D** The *D*/*k* values of the BraTS tumors (*n = 91*) grouped by gene-expression subtyping from microarray data of 202 TCGA glioma samples^24^. **E** The *D*/*k* values of BraTS tumors (*n = 81*) grouped by RNA-seq expression subtypes from 667 TCGA glioma samples^25^. **F** The *D*/*k* values of clusters of BraTS tumors (*n = 22*) based on random forest classification of DNA methylation profiles of 324 glioma samples^25^.

### Glycosylation Pathway associations identified by Elastic Net Regression and overrepresentation analysis

We performed a coarse hyperparameter sweep followed by a fine hyperparameter sweep over four Elastic Net hyperparameters: the variance threshold, the most correlated/anti-correlated subset size, the weighting between LASSO and ridge regression, and the weighting between these penalties and least-squares (Figure S4). Using the hyperparameter set that produced the greatest r-squared value with LOOCV, we plotted the predicted *D*/*k* for all left-out patients and found a significant correlation (Pearson *r = 0.42, p = 3.3X10^−5^*). We constructed a histogram of the frequency with which each gene was given a nonzero weight by Elastic Net over all 91 LOOCV runs (Figure 6B). We next ran Elastic Net regression with the full dataset, and highlighted the nonzero weighted genes in the histogram (*n = 98*, Figure 6B, orange bars). Given the strong agreement between the most frequently selected genes in the LOOCV runs and the full dataset run (Figure 6B), we plotted the expression of these genes with patients sorted by *D*/*k* (Figure S5).

**Figure 6:**
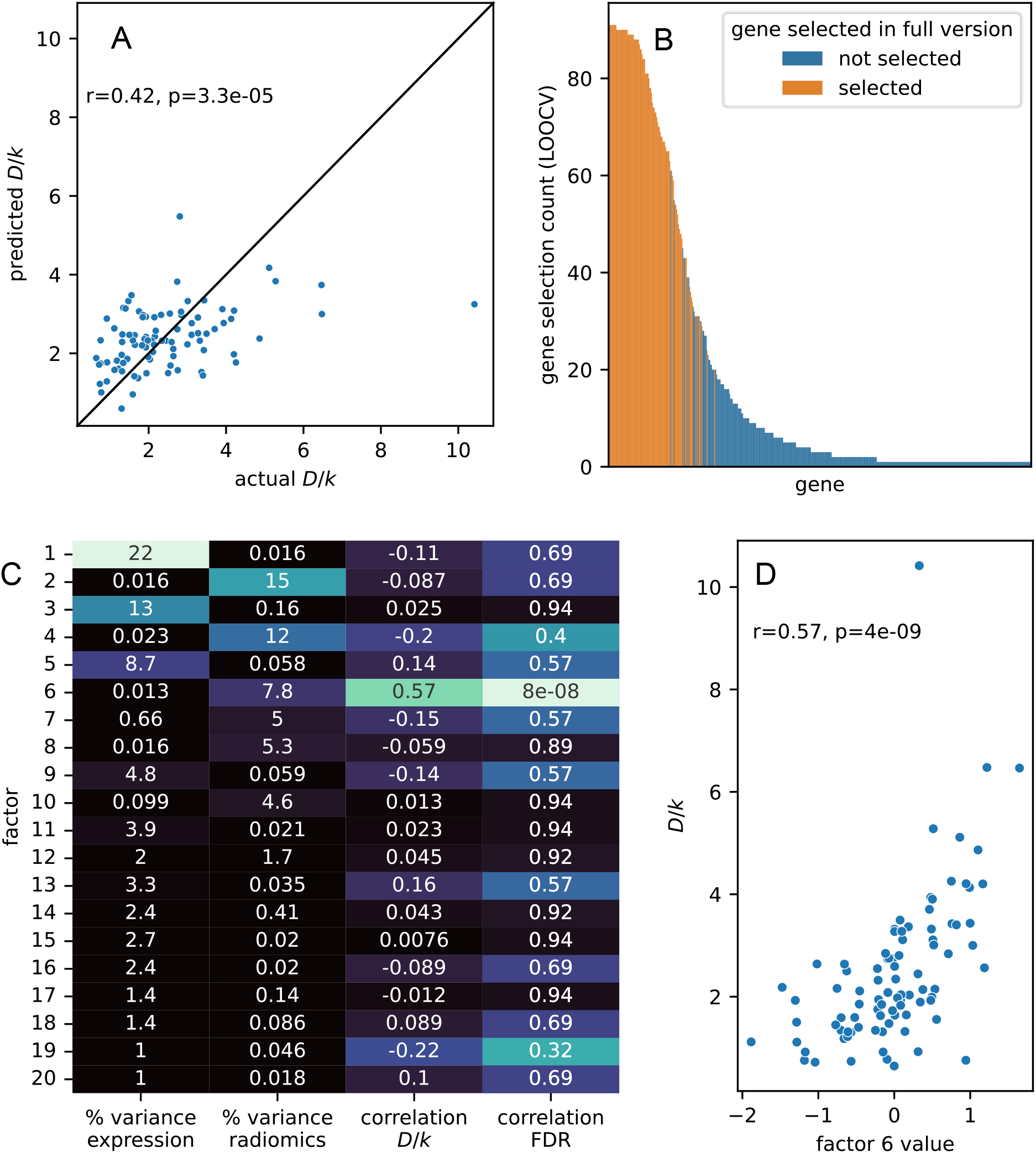
Multi-omic and transcriptomic correlates of *D*/*k*: **A** Scatter plot of LOOCV predicted *D*/*k* vs actual *D*/*k* for each left-out patient. **B** Histogram showing frequency that the same gene was chosen by Elastic Net during iterations of LOOCV. Color indicates whether the gene was also selected by Elastic Net when all patients were included in training. **C** Heatmap showing % variance explained by each MOFA factor and how much each factor correlates with *D*/*k*. **D** Scatter plot showing correlation between factor 6 score and *D*/*k*.

Finally, we used overrepresentation analysis (ORA) to identify biological pathways significantly enriched within the genes selected by Elastic Net regression using the full dataset (Figure 6B). We performed ORA using the reactome and hallmarks gene set libraries^48,49^. The Reactome gene sets “Asparagine N-linked Glycosylation” and “Synthesis of UDP N Acetyl Glucosamine” were the only sets with significant enrichment after Bonferroni correction, with adjusted p-values of *0.0123* and *0.0368*, respectively. None of the msigdb hallmark gene sets were significantly enriched after Bonferroni correction (Table S2).

### Multi-omics factor analysis finds radiomic factor correlated with D/k

Applying MOFA to the combined radiomic and microarray data produced 20 factors that describe variance across both datasets (Figure 6C). The factor 6 significantly correlates with *D*/*k* (r = 0.57, p = 4X10⁻⁹; Figure 6D). The features with the highest magnitude weight for factor 6 are listed in Table 2.

**Table 1:** Results from ORA done on genes chosen by Elastic Net run on all patients.

| Term | Overlap | P-value | Adjusted P-value | Odds Ratio | Combined Score | Genes |
|---|---|---|---|---|---|---|
| REACTOME_ASPARAGINE_N_LINKED_GLYCOSYLATION | 7/246 | 5.62E-05 | 0.0123 | 8.664 | 84.79 | AMDHD2;RENBP;RPN2;LMAN2L;OS9;RPN1;GRIA1 |
| REACTOME_SYNTHESIS_OF_UDP_N_ACETYL_GLUCOSAMINE | 2/7 | 3.36E-04 | 0.0368 | 114.8 | 917.9 | RENBP;AMDHD2 |
| REACTOME_MATURATION_OF_SARS_COV_2_SPIKE_PROTEIN | 2/28 | 0.0057 | 0.2327 | 23.78 | 122.7 | RPN2;RPN1 |
| REACTOME_DIFFERENTIATION_OF_KERATINOCYTES_IN_INTERFOLLICULAR_EPIDERMIS_IN_MAMMALIAN_SKIN | 2/29 | 0.0061 | 0.2327 | 22.91 | 116.7 | TFRC;EGF |
| REACTOME_CARGO_CONCENTRATION_IN_THE_ER | 2/31 | 0.007 | 0.2327 | 21.35 | 106 | LMAN2L;GRIA1 |

**Table 2:** Top 10 features with greatest magnitude weight from MOFA factor 6. “Runs” refer to neighboring voxels along a line that have the same intensity value. ^50^.

| Feature | Description | Weight |
|---|---|---|
| TEXTURE_GLRLM_ED_T2_RLN | Heterogeneity in run sizes in edema region on T2 | -1.27 |
| TEXTURE_GLRLM_ED_T2_SRE | Prevalence of short runs in edema region on T2 | -1.27 |
| TEXTURE_GLSZM_ED_FLAIR_ZP | Proportion of contiguous intensity zones relative to the number of voxels in edema region on FLAIR | -1.25 |
| TEXTURE_GLRLM_ED_T2_RP | Ratio of number of runs to number of voxels in edema region on T2 | -1.25 |
| TEXTURE_GLRLM_ED_T2_LRE | Prevalence of long runs in edema region on T2 | 1.20 |
| TEXTURE_GLRLM_ED_FLAIR_RLN | Heterogeneity in run sizes in edema region on FLAIR | -1.20 |
| TEXTURE_GLRLM_ED_FLAIR_SRE | Prevalence of short runs in edema region on FLAIR | -1.19 |
| TEXTURE_GLSZM_ED_T2_ZP | Proportion of contiguous intensity zones relative to the number of voxels in edema region on T2 | -1.19 |
| TEXTURE_GLRLM_ED_FLAIR_RP | Ratio of number of runs to number of voxels in edema region on FLAIR | -1.18 |
| TEXTURE_GLCM_ED_T2_Correlation | Similarity of gray-level values between neighboring voxels in edema region on T2 | 1.16 |

## Discussion

### D/k pipeline confirmed with synthetic density distributions and segmentations

The synthetic density distribution experiments demonstrated that the D/k pipeline accurately recovers known D/k values, with estimates generally within 1-3% of the true values. Holding either k or D constant produced similar estimates, but we chose to hold k constant because this resulted in more consistent simulation durations across D/k values.

Adding noise to the line profiles causes a greater deviation when *D*/*k* is low (0.1) than when *D*/*k* is high (10.0). One likely explanation is that the gradient region is smaller when *D*/*k* is low. In this regime, tumor density drops off steeply, leaving fewer voxels over which noise can average out and making the estimated gradient more sensitive to noise.

From fitting segmented density distributions, we found that fits were consistent despite the decrease in detail caused by the segmentations and testing different tumor sizes. These test estimates had a consistent bias but were highly correlated with the true D/k values (PCC = 0.996).

### D/k is correlated with age and overall survival

We evaluated whether patient-specific *D*/*k* estimates were associated with clinical measurements from the BraTS cohort. We found that patient groups stratified by *D*/*k* exhibited a significant difference in overall survival based on Kaplan-Meier analysis (log-rank p = 9.89 × 10⁻³). This indicates that higher *D*/*k* is associated with poorer overall survival. We also observed that *D*/*k* correlates with age (p = 0.04) and negatively correlates with functional performance (KPS score, p = 0.06). The association between D/k and overall survival may not be independent of age, as the negative correlation between age and overall survival in GBM is well documented^51,52^. These results were also consistent when we applied the Swanson et al. *D*/*k* method^11^ to the dataset, (Figure 5, Figure S3), supporting the consistency of these associations with a different *D*/*k* estimation method. Our *D*/*k* values and Swanson-method *D*/*ρ* values for all patients are also highly correlated with each other, which is expected given that both methods quantify the ratio of migration to proliferation.

Seeing the strong correlations between *D*/*k* and other clinical data suggests that it could be a meaningful descriptor of tumor aggressiveness. If this is the case, *D*/*k* could potentially inform surgical margin decisions, where patients with greater *D*/*k* would be expected to have more cells outside of the detectable margin than patients with lower *D*/*k*^8,53,54^. This could help personalize the margins of surgery and radiotherapy beyond the detectable tumor to maximize remission time while limiting damage to healthy tissue. In addition, lower *k* has been associated with reduced radiotherapy effectiveness^53^, suggesting that patient-specific estimation of *D* and *k* could further improve radiotherapy personalization.

### Mesenchymal tumors tend to exhibit increased D/k

Of the 102 patients in the BraTS 2017 dataset, 91 had microarray data available as part of the TCGA, which provided the opportunity to identify transcriptional patterns that are associated with *D*/*k*. We observed a consistent (though not always significant) trend when we classify the patients by molecular subtypes^24,25^, where the mesenchymal subtype tends to show a greater *D*/*k* than the other subtypes. This follows the expectation that mesenchymal tumors are more migratory than the other subtypes. Additionally, patients with mesenchymal-like tumors tend to be older^22^, which connects to the correlation between *D*/*k* and age (Figure 5B). Mesenchymal tumors are also more resistant to radiotherapy and chemotherapy^22^, which could explain the poorer overall survival of elevated *D*/*k* patients.

### Glycosylation pathways associated with increased D/k

Overrepresentation analysis of the genes identified by Elastic Net revealed significant enrichment of two Reactome pathways: Asparagine N-linked glycosylation (adjusted p = 0.0123) and Synthesis of UDP-N-acetylglucosamine (adjusted p = 0.0368). These pathways together associate N-linked glycosylation-related processes with interpatient variation in *D*/*k*, as UDP-GlcNAc (UDP-N-acetylglucosamine) is an indispensable component of N-linked glycosylation^55–57^ and is also associated with increased migration in gliomas^60^.

Glycosylation is the enzyme catalyzed addition of sugar complexes to proteins, and is a particularly prevalent modification of cell-surface proteins^56^. Increased N-linked glycosylation has been associated with increased migration through degrading cadherin-catenin complexes which decreases cell-cell adhesion^55,58,59^, and conversely decreased N-linked glycosylation has been associated with strengthening adhesion and thus limiting migration^57^. Increased N-linked glycosylation also affects turnover of integrin binding to the ECM, which further facilitates migration^57^. This makes glycosylation a plausible contributor to the cell-cell adhesion association with *D*/*k* reported by Metz et al ^10^. Increased transcription related to glycosylation has also been previously associated with mesenchymal subtype in GBM^60^ and epithelial to mesenchymal transition in general^56,61^.

### Radiomic texture of the non-enhancing region may provide an imaging biomarker of D/k

Applying Multi-omic Factor Analysis (MOFA)^26^ to the combined radiomic and microarray data, we found that the factors tended to explain the variance in either microarray or radiomic features but not both, indicating that the dominant sources of variation identified by MOFA were largely modality-specific in this dataset.

We identified one factor that is highly correlated with *D*/*k,* and upon examining the highest weighted features, they were all radiomic features in the non-enhancing tumor on either *T_2_* or FLAIR. The consistent trend indicated by the features and the signs of their associated weights is that coarser, more homogeneous texture in the non-enhancing region is associated with high D/k, whereas finer, more heterogeneous texture is associated with low D/k. This makes intuitive sense, as increased random migration would be expected to smooth gradients in tumor density, which would cause more uniformity in intensity values on imaging. This suggests that the level of uniformity of the texture of edema on *T_2_* and FLAIR MRI scans could be an imaging surrogate of the *D*/*k* phenotype.

### Limitations and future directions

In constructing the tumor density distributions from imaging, we made the simplifying assumption that the different segmented regions had constant tumor density. This allowed us to ignore intratumor heterogeneities (e.g., necrosis), which would have been difficult to incorporate into the model and fit parameters with only a single MRI per patient. However, this constant density assumption is an oversimplification of the true tumor density gradient found in GBM. Although ADC scans could have helped determine this patient-specific density gradient^32–34^, datasets containing both advanced imaging and matched molecular profiling are relatively rare. We therefore utilized the BraTS dataset because it provides one of the largest cohorts with both tumor segmentations and genomic data. We also assumed that the normal-appearing tissue has no tumor cells whatsoever. This is not the case in reality as GBM recurrence often happens even with wide surgical margins^62^, though the tumor cell density in this region tends to be very low^63^.

We also did not directly incorporate many known anatomical features that impact tumor cell migration, like differential migration rates through grey and white matter regions^64^, the boundaries of the brain itself, or the impact of tissue compression^65^. Incorporating these details could result in estimating tumor migration more accurately. However, these additional assumptions would dramatically increase the model complexity, challenging parameter estimation, and would almost certainly require additional imaging time points.

Although tumor imaging allows for the assessment of the entire tumor density distribution in 3D, the associated microarray data only characterizes the tumor transcriptome at one biopsy location. Given intratumoral heterogeneity^66^ and the importance of context to cell phenotype^67^, this may have been an unaccountable covariate in our analyses. Microarray analysis also provides a less extensive profile than RNA-seq^68^ which may have also impacted our results. It would thus be useful to confirm these results with an RNA-seq dataset, ideally from consistent biopsy locations within the tumor or from recent spatial transcriptomics studies^69^.

Furthermore, glycosylation itself is a very broad category of protein modifications that is estimated to modify over half of the human proteome^55,57^. Although the category we identified, N-linked glycosylation, aligns with the GBM literature^55,58,59,70^, the specific proteins that are being modified cannot be directly investigated with transcription data alone. Advanced multi-omic datasets such as The Clinical Proteomic Tumor Analysis Consortium (CPTAC) GBM dataset^71^ and associated glycoproteomics data^72^, are needed to provide direct quantification of the glycoproteins associated with variation in *D*/*k*.

Another finding that warrants further investigation is the association between *D*/*k* and the uniformity of the non-enhancing region on FLAIR and *T_2_*-weighted imaging. If this association is validated, it could serve as a spatially localized surrogate of *D*/*k* phenotype, which could further improve patient-specific predictions of tumor invasive margins beyond the global *D*/*k* metrics presented. This could be tested with longitudinal tumor imaging data so that predicted differences in migration and growth from local radiomic features could be verified over time. Radiomic features of the non-enhancing region have shown some predictive capability of recurrence location in GBM, providing evidence that supports this idea^73^.

## Conclusions

We validated our *D*/*k* pipeline by accurately recovering *D*/*k* from a 3D PDE-derived density distribution, and found strong correlation when these distributions were grown to various sizes and converted to segmentations prior to fitting. By analyzing associations between patient *D*/*k* values and clinical variables, we found that high *D*/*k* is associated with poor survival outcomes, older age, and shows a trend towards lower KPS. Analysis of microarray-derived subtypes suggests that the mesenchymal subtype is associated with higher *D*/*k*. Our Elastic Net regression based selection of predictive genes followed by overrepresentation analysis associates N-linked glycosylation-related pathways with interpatient variation in *D*/*k*, consistent with its known role in promoting tumor migration. Finally, our Multi-Omic Factor Analysis (MOFA) of microarray expression and radiomic features suggests that homogeneous regions on FLAIR and *T_2_*-weighted MRI are associated with increased *D*/*k.* Collectively, these findings support *D*/*k* derived from a single pre-treatment MRI as a promising non-invasive biomarker associated with patient survival that also captures biologically meaningful differences in molecular phenotype.

## Acknowledgements

We thank the National Institutes of Health for support through R01CA260003 and the National Science Foundation for support through DMS-2436499. TEY is a CPRIT Scholar in Cancer Research. BT and BA received funding from the National Cancer Institute of the National Institutes of Health under award number R01CA270210. We thank Param Hansa Philanthropies for supporting the UT Austin Center for Computational Oncology.

## Data Availability Statement

The MRI data, tumor segmentations, and clinical information analyzed in this study were obtained from the BraTS 2017 dataset, available through the Center for Biomedical Image Computing and Analytics at the University of Pennsylvania (https://www.med.upenn.edu/sbia/brats2017/registration.html). The corresponding TCGA-GBM imaging data, segmentation labels, and radiomic features are available through The Cancer Imaging Archive (https://doi.org/10.7937/K9/TCIA.2017.KLXWJJ1Q).

Gene expression data were obtained from The Cancer Genome Atlas (TCGA) glioblastoma cohort through the Broad Institute’s FireBrowse portal (https://firebrowse.org/; https://gdac.broadinstitute.org/).

The code implementing the D/k estimation method, together with documentation and reproducible synthetic examples, is available at https://github.com/OncologyModelingGroup/DkMethod/. The repository also includes patient-level D/k estimates for 102 cases, coefficients for 98 selected genes, and code for downstream clinical and molecular analyses. The original MRI, clinical, and molecular datasets are not redistributed. Researchers can access the original datasets through the sources listed above and use the provided code to estimate D/k from patient imaging data.

## Supplemental

**Table S1:**
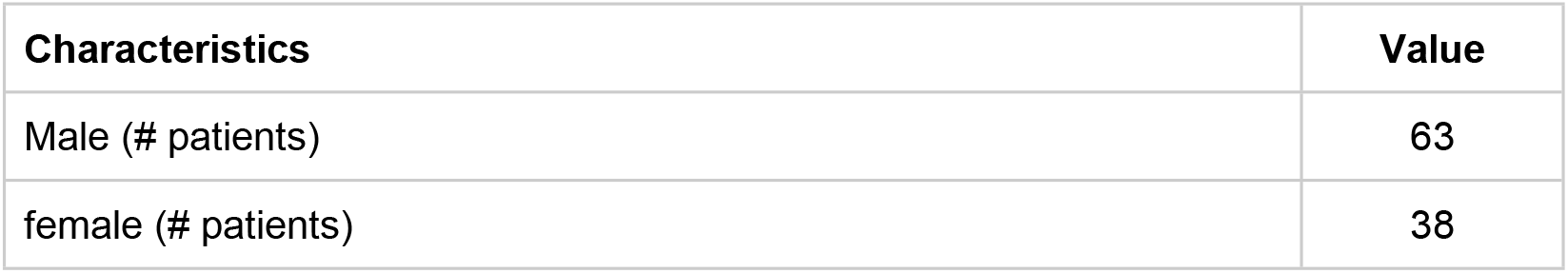
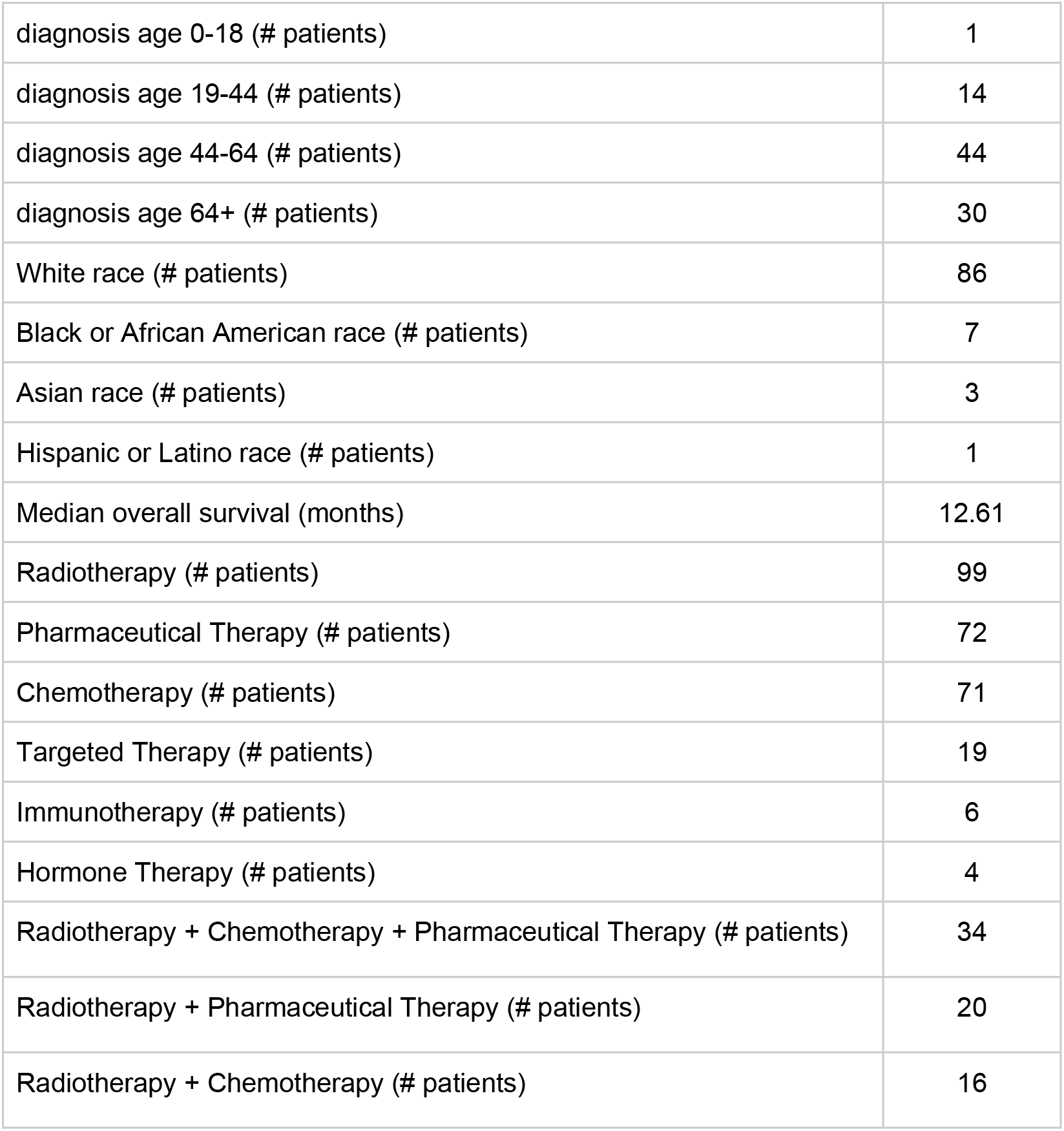
BraTS 2017 patient demographic information. All patients underwent surgery.

**Figure S1:**
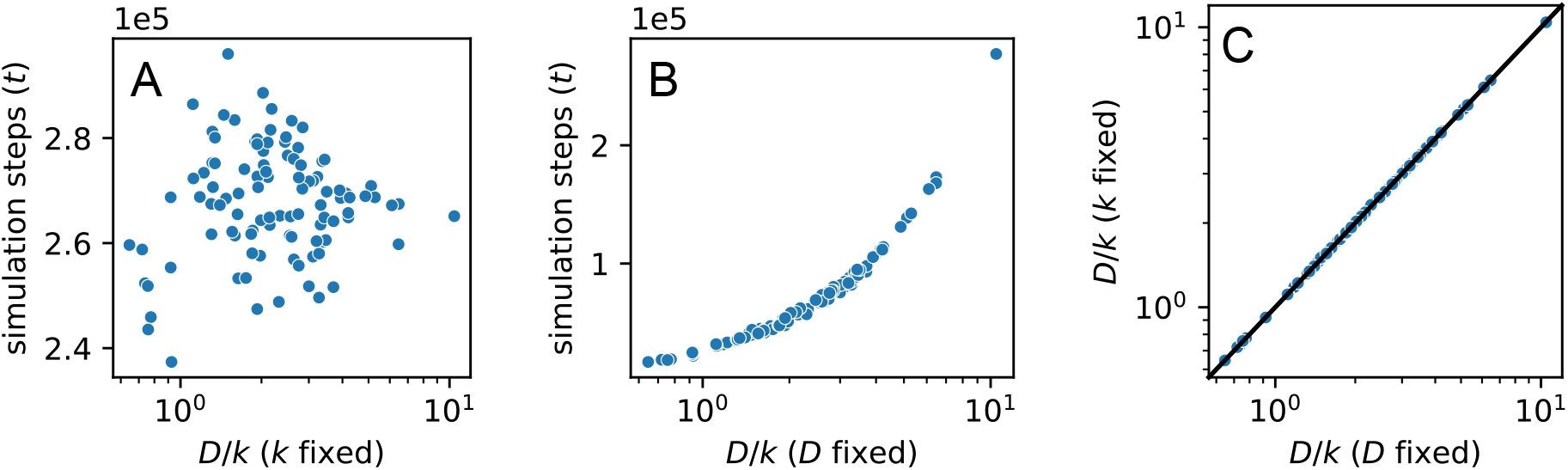
**A, B** *D*/*k* ratios and optimal numbers of simulation steps calculated for the 102 patients from the BraTS 2017 dataset, with *k* fixed (panel A) and *D* fixed (panel B). **C** *D*/*k* ratios with *k* fixed plotted against *D*/*k* ratios with *D* fixed. CCC = 1.000

### Starting the synthetic tumor near the domain boundary distorts the inferred D/k

We also determined the impact of initiating the synthetic tumor near the edge of the computational domain on the ability to accurately estimate *D*/*k* (Figure S2A). For this test, we assessed two different methods for finding the tumor initial position: the center of mass of the enhancing region (Figure S2B) and the intersection point of the two longest perpendicular diameters within the enhancing region (Figure S2C). The center of mass method showed reduced correlation with the synthetic *D*/*k* values when the tumor initial position was within 10 mm of the boundary compared to 20 mm (PCC = 0.927 versus 0.995), and the longest diameters method maintained high correlation regardless of boundary proximity (PCC = 0.997 versus 0.999).

When applied to the patient data, however, we observed that the longest diameters method tended to generate larger *D*/*k* values relative to the center of mass method. This is the opposite of the correction that we observed with the synthetic data (Figure S2D, E). This discrepancy is likely due to the longest perpendicular diameters method selecting tumor initial positions that are farther from the center of the enhancing region in asymmetrical tumors (Figure S2F). We thus chose the center of mass method because of its simplicity and consistency when applied to the patient data.

**Figure S2:**
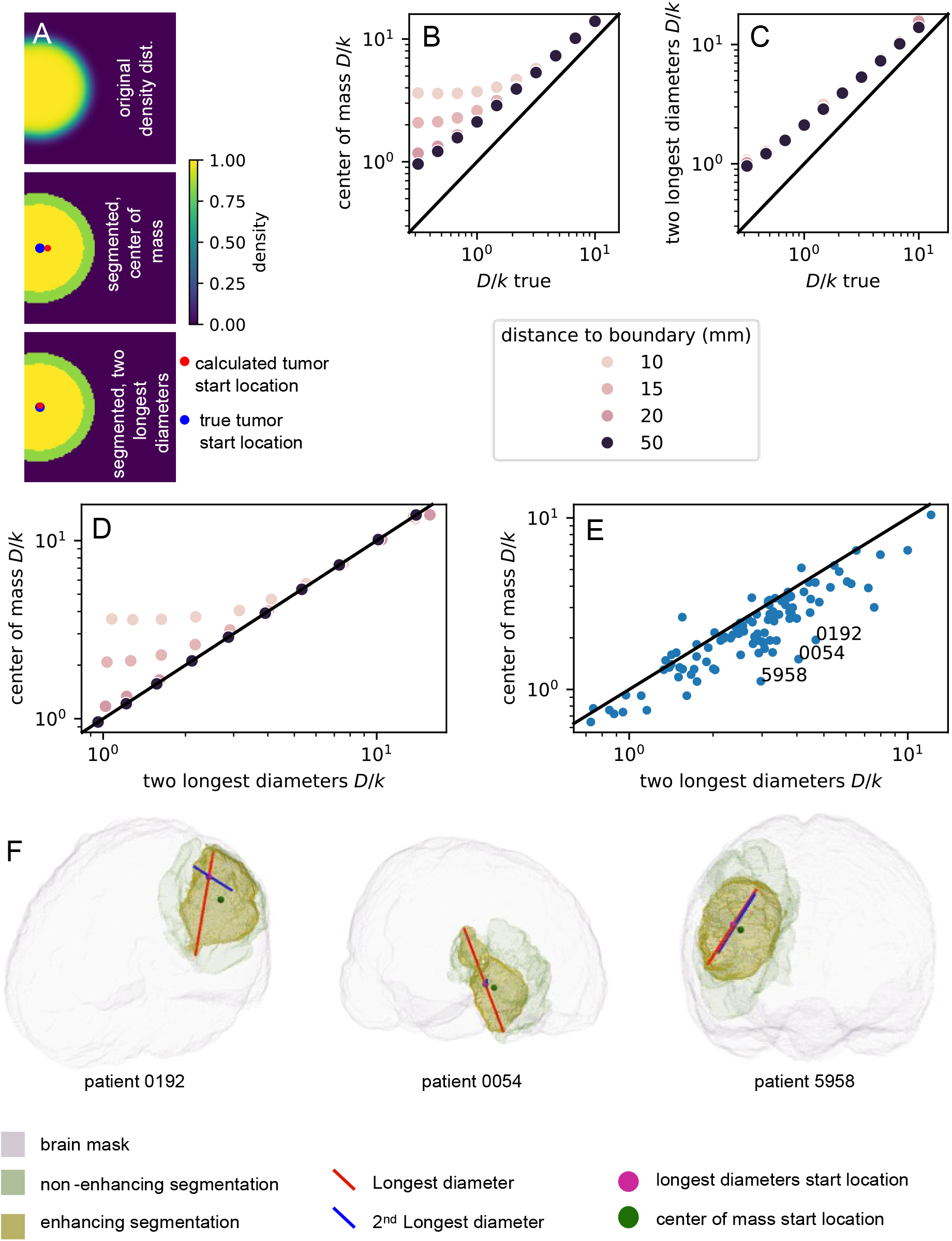
Boundary proximity effects and tumor initial position estimation methods **A** Top: 2D slice of 3D model run where tumor initial position is near the edge of the domain, middle and bottom: segmented tumor with initial position calculated via center of mass method and longest diameters method. **B, C** *D*/*k* calculated via center of mass method and longest diameters method versus synthetic data *D*/*k* (*D*/*k_true_*) for different distances of the tumor initial position from the boundary. **D, E** *D*/*k* calculated from the two longest diameters method versus the center of mass method for the **D** Synthetic data and **E** 102 patients from the BraTS 2017 dataset. **F** Three patients with particularly large discrepancies in calculated *D*/*k* are plotted in 3D to demonstrate the difference in tumor initial position between the longest diameters method and the center of mass method.

**Figure S3:**
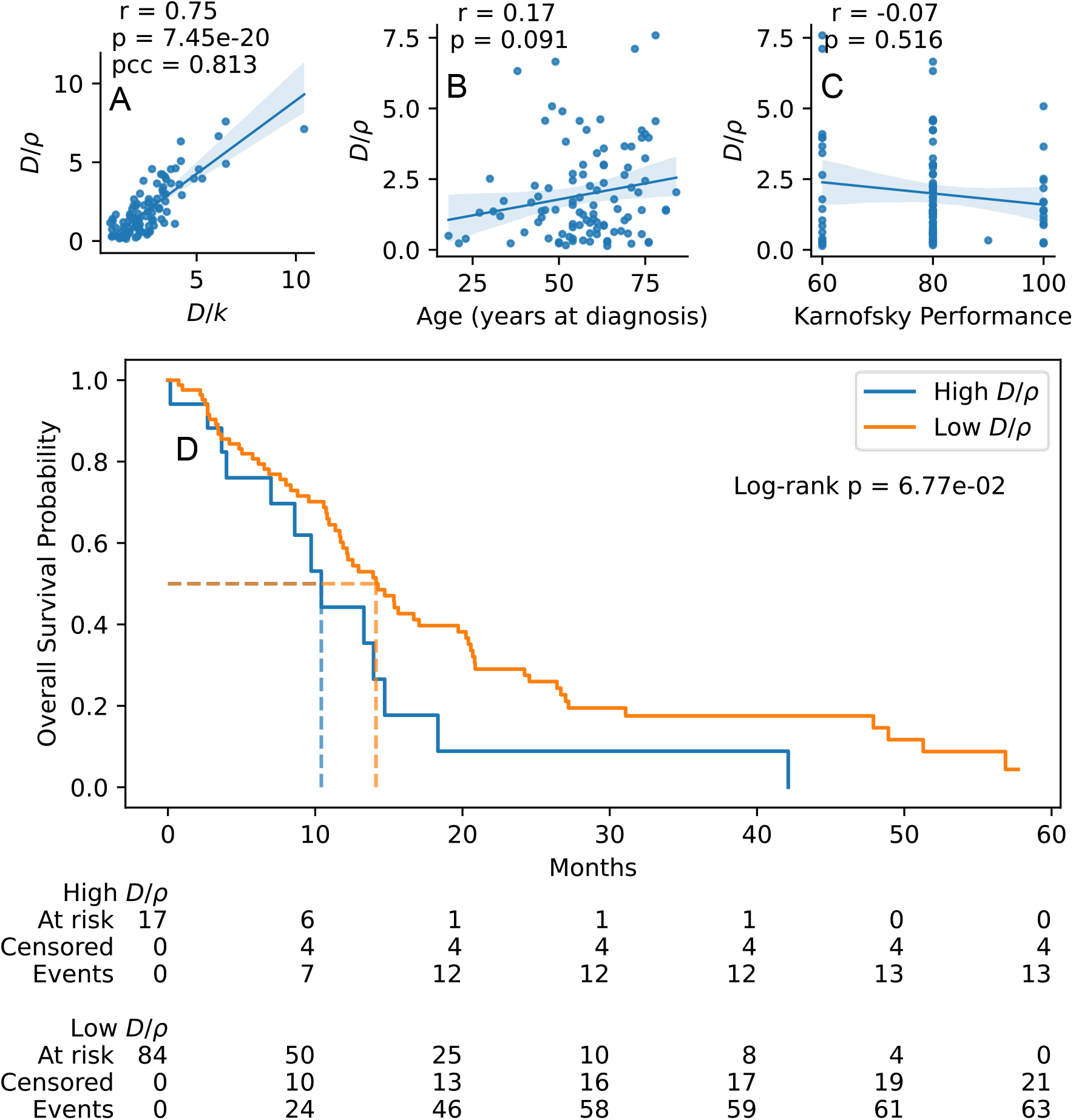
Comparison of *D*/*k* with the Swanson *D*/*⍴* method and clinical outcomes. **A** Correlation between *D*/*k* and Swanson *D*/*⍴*^11^ method applied to BraTS patient data. *D*/*⍴* found using Equation (1) from Tropeano et al.^12^. **B, C** Correlation of patient age versus *D*/*⍴* and Karnofsky Performance Score versus *D*/*⍴*. Spearman correlation was used to calculate *r* and p values. **D** Optimal cut point between 10th and 90th percentiles of the patient data ordered by *D*/*⍴* was determined via log-rank test to separate patients into Low *D*/*⍴* and High *D*/*⍴* groups. Kaplan-Meier curves of optimally cut groups are shown with associated log-rank *p*-value.

**Figure S4:**
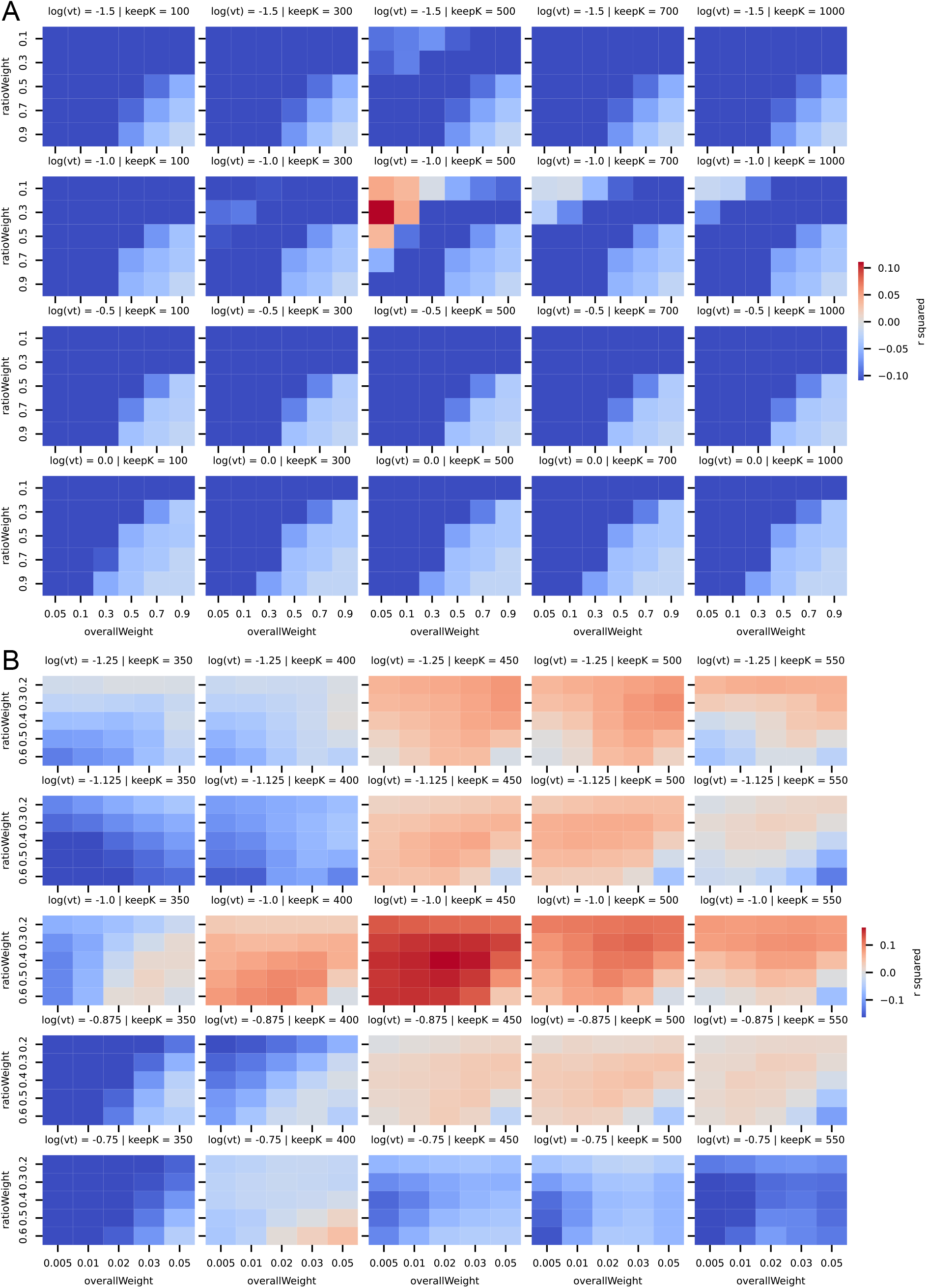
**A, B Elastic Net Regression:** hyper parameter tuning performance. Within heatmaps: rows indicate weighting of ratio between ridge regression and LASSO regression, columns indicate weighting of combination of ridge and LASSO regression penalty versus ordinary least-squares regression. Across heatmaps: rows indicate gene microarray variance threshold as part of pre-processing, columns indicate the number of genes most correlated with *D*/*k* that are subsequently fed into Elastic Net.

**Figure S5:**
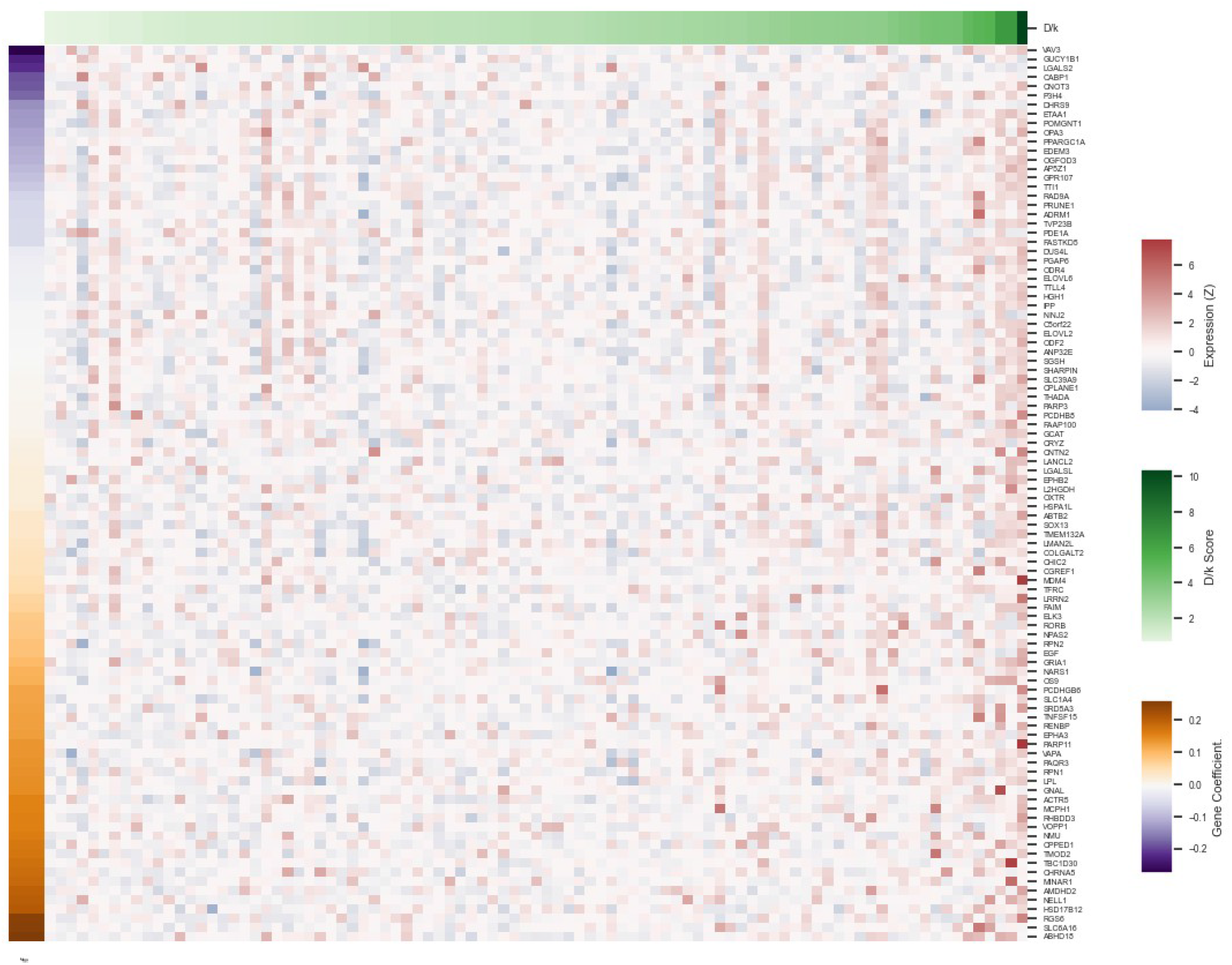
Microarray expression of genes chosen by Elastic Net regression model trained on the full patient dataset. The top annotation bar indicates the estimated *D*/*k* for each patient. Left annotation bar indicates the linear model coefficients.

### Gene set analysis results

We performed overrepresentation and GSVA^74^ analysis for Reactome pathways and the most positively correlated Reactome pathways with *D*/*k* are listed in Table S3. Many Reactome pathways were significantly correlated with before multiple test correction, they were not significant after.

In overrepresentation analysis, the closest was to the UV_RESPONSE_UP gene set and the MTORC1_SIGNALING gene set. Upregulation of UV response genes could be linked to decreased drug response in elevated-*D*/*k* patients, as the genes in this pathway are associated with DNA repair, which could also dampen response to chemotherapy and radiotherapy. The mTOR signaling reflects multiple cellular processes including proliferation and migration, suggesting that elevated mTOR activity corresponds to more advanced disease.

**Table S2:** Results from ORA done on genes chosen by Elastic Net regression run on all patients (msigdb database).

| Term | Overlap | P-value | Adjusted P-value | Odds Ratio | Combined Score | Genes |
|---|---|---|---|---|---|---|
| HALLMARK_UV_RESPONSE_UP | 3/148 | 0.0222 | 0.3577 | 6.1314 | 23.3517 | RPN1;CHRNA5;TFRC |
| HALLMARK_MTORC1_SIGNALING | 3/186 | 0.0397 | 0.3577 | 4.8461 | 15.63 | SLC1A4;RPN1;TFRC |
| HALLMARK_ANGIOGENESIS | 1/36 | 0.1367 | 0.5342 | 10.4218 | 20.7418 | LPL |
| HALLMARK_ESTROGEN_RESPONSE_LATE | 2/189 | 0.1792 | 0.5342 | 3.3152 | 5.6998 | SLC1A4;NMU |

**Table S3:** Reactome pathways whose GSVA scores are most correlated with the estimated *D*/k *.

| **Pathway** | **Spearman_Rho** | **P_Value** |
|---|---|---|
| HS-GAG Degradation | 0.3178 | 0.0021 |
| Diseases of Carbohydrate Metabolism | 0.3011 | 0.0037 |
| Diseases of Metabolism | 0.2982 | 0.0041 |
| Diseases of Glycosylation | 0.2743 | 0.0085 |
| Defective EXT2 Causes Exostoses 2 | 0.2726 | 0.0089 |
| Defective EXT1 Causes Exostoses 1, TRPS2 and CHDS | 0.2726 | 0.0089 |
| HS-GAG Biosynthesis | 0.2606 | 0.0126 |
| Mucopolysaccharidoses | 0.2552 | 0.0146 |
| Keratan Sulfate Degradation | 0.2515 | 0.0162 |
| Heparan Sulfate Heparin (HS-GAG) Metabolism | 0.2493 | 0.0172 |

\*No significant results after FDR correction.

## Notes

### Competing Interest Statement

The authors have declared no competing interest.

